# Evaluation of the possible association of Body Mass Index and four metabolic gene polymorphisms with longevity in an Italian cohort: a role for *APOE, eNOS*, and *FTO* gene polymorphisms

**DOI:** 10.1101/608745

**Authors:** Alfredo Santovito, Gabriella Galli, Stefano Ruberto

## Abstract

**Background:** longevity is considered the result of interactions between environmental and genetic factors.

**Aim:** we investigated the possible association of body mass index and the frequencies of *APOE, ACE, eNOS*, and *FTO* gene polymorphisms with longevity.

**Subjects and Method:** 1,100 healthy volunteers aged 10-100 were recruited. We genotyped subjects for *APOE, ACE, eNOS*, and *FTO* gene polymorphisms. Data about height and weight were also collected. The sample was split in four age groups: 1-24, 25-49, 50-85 and 86-100.

**Results:** significant differences were found in BMI values between age groups, with exception of 1-24 with respect to 86-100. A significant decrease of the *APO E4, eNOS 393* and *FTO A* and allele frequencies was observed in the 86-100 age group with respect to younger groups. For *ACE* gene, no significant differences were found in the allele frequencies between groups. A similar trend was also observed subdividing the sample in two main age groups: 1-85 and 86-100.

**Conclusion:** this study provides evidences for a role of *APOE, eNOS*, and *FTO* gene polymorphisms in longevity. It has been estimated that the number of centenarians worldwide will double each decade until 2100, making population data about gene polymorphisms relevant for further studies about longevity.

## 1. INTRODUCTION

Longevity is generally considered as the result of interactions between environmental and genetic factors. Among the environmental factors, life-style and the nutritional status seem to play an important role in the incidence of many age-related diseases. For example, high levels of adiposity, typical of individuals with a body mass index (BMI) more than 30 kg/m^2^, were linked to abnormal glucose metabolism and increased incidence of cardiovascular diseases (CVDs), cancer and neurodegenerative diseases, with consequently higher mortality rate (Srikanthan and Karlamangla, 2014).

Moreover, since CVDs are the main causes of death worldwide (WHO, 2011), polymorphisms in genes involved in these types of diseases, as well as those involved in vascular dysfunction and obesity, represent excellent candidates for longevity.

Apolipoprotein E (*APOE*) gene is one of the most consistently associated with human longevity (Fuku et al., 2017). The glycoprotein encoded by this gene plays a fundamental role in the lipid metabolism, mediating lipoprotein binding to the LDL and lipoprotein remnant receptors. Defects in this protein could diminish its ability to bind to the receptors, which then leads to an elevated blood cholesterol level and, consequently, to increased risk of atherosclerosis and CVDs (Marrzoq et al., 2011). *APOE* gene is polymorphic, with three common alleles *E2, E3*, and *E4*, that give rise to six different genotypes *E2/2, E2/3, E2/4, E3/3, E4/3*, and *E4/4* (Marrzoq et al., 2011). The *E4* allele is the most common *APOE* allele in all human populations and it is associated with higher total cholesterol levels and increased risk of metabolic and neurodegenerative diseases, whereas *E2* and *E3* alleles were associated with lower cholesterol levels and with a protective role against Alzheimer’s disease (de-Almada et al., 2012; Marrzoq et al., 2011).

Another gene associated with CVDs is the *ACE* gene, that encodes an enzyme that catalyzes the conversion of angiotensin I to angiotensin II, a potent vasoconstrictor. In human populations this gene is polymorphic due to the presence (insertion, allele I) or absence (deletion, allele D) of an *Alu* sequence of 287-bp in intron 16. In some studies, individuals with Alzheimer’s disease have been reported to show an increased frequency of I allele, while homozygosity of the *ACE* D allele has been associated with an increased risk for CVDs and hypertension (Garatachea et al., 2013).

Endothelial nitric oxide, synthesized by nitric oxide synthases gene (*eNOS*), plays a crucial role in the regulation of vascular tone. It diffuses from the endothelium to vascular smooth muscle cells, causing vascular relaxation and inhibiting the cellular proliferation. During aging, there is a progressive reduction of NO production, whereas the oxidative stress increases as a consequence of the absence of a compensatory enhancement of antioxidant defences. The result of this reduction of the available NO, is the alteration of the vascular homeostasis with possible increased risk for hypertension, atherosclerosis, thrombosis, and stroke (Puca et al., 2012). Several polymorphisms were reported in the promoter region, exons and introns of this gene. An important one is represented by a 27-base pair core consensus VNTR (variable number of tandem repeats) polymorphism in the intron 4: the common wild-type ‘b-allele’ (*eNOS4b* allele) has five tandem 27-bp repeats, while the rare ‘a-allele’ (*eNOS4a* allele) only four repeats. In particular, the *eNOS* 4a allele was found to be an independent risk factor of myocardial infarction, whereas the *eNOS* 4a/4a homozygote was identified as a smoking dependent risk factor of myocardial infarction (Park et al., 2004).

Finally, fat mass and obesity-associated (*FTO*) gene is another CVDs risk candidate gene, also associated to obesity and BMI. The common single-nucleotide polymorphism, rs9939609, located in the first intron of the FTO gene is one of the most important variants that have been examined by many genome wide association studies, with the allele A strongly associated with obesity phenotype, increased BMI values, CAD and metabolic syndrome (Qi et al., 2014; Reitz et al., 2012).

The main objectives of this study were to investigate the possible association of BMI, of some metabolic (*APOE, ACE, eNOS*, and *FTO*) gene polymorphisms and longevity in a northern Italy cohort. The hypothesis is that some anthropometric parameters, such as the BMI, and some genetic variants could be differentially represented in long-lived individuals with respect younger subjects.

## 2. MATERIALS AND METHODS

### 2.1 Study population

The study was conducted on a cohort of 1,100 healthy Italian subjects (aged 10-100 years; 546 males and 554 females) of Caucasian origin from Northern Italy. All the subjects were randomly chosen healthy volunteers, received detailed information about the study, were anonymously identified by a numeric code and gave their informed consent prior the analyses. Inclusion criteria for the control group were being a man or woman with no history of stroke, cardiovascular disease, diabetes, neurodegenerative and cancer diseases. The research protocol was approved by the local ethics committee and was performed in accordance with the ethical standards laid down in the 2013 Declaration of Helsinki.

In Italy, in 2017, the average life expectancy at birth was 80.6 years for men and 84.9 years for women (ISTAT, 2017a). Moreover, the fertility rate for women belonging to the 20-49 age group was about 90%, with an average age of mothers and fathers at the birth of their first child of 31.8 and 35.3 years, respectively (ISTAT, 2017a,b). For these reasons, in order to evaluate the possible association of some alleles with longevity, we decided to divide our sample into 4 age groups: 1-24, that include subjects in pre-reproductive phase; 25-49, that included subjects in reproductive phase; 50-85, representing the group of subjects in post-reproductive phase; and 86-100 that represent the group with long-lived individuals.

### 2.2 DNA extraction and Genotyping

Peripheral blood samples (5-10 ml obtained by venipuncture) were collected in heparinized vacutainers and stored at −20°C. DNA extraction was conducted using a Chelex solution, according to Santovito et al. (2017). Gene polymorphisms were determined by PCR and RFLP methodologies, using primers and melting temperature as described in Table 1. PCR reactions were performed in a 25 μL volume containing about 10 ng DNA (template), with a final concentration of 1X Reaction Buffer, 1.5 mM of MgCl_2_, 5% of DMSO, 250 μM of dNTPs, 0.5 μM of each primer, and 1 U/sample of Taq DNA polymerase (Fischer, U.S.). Cycles were set as follows: 35 cycles, 1 min at 95°C, 1 min at 56-65°C depending on the primer sequence (Table 1), 1 min at 72°C, and a final extension step 10 min at 72 °C. Amplification products were detected by ethidium bromide staining after 4% metaphor agarose gel electrophoresis. The expected PCR and/or RFLP product size for each gene polymorphism analysed were showed in Table 1.

**Table 1.**
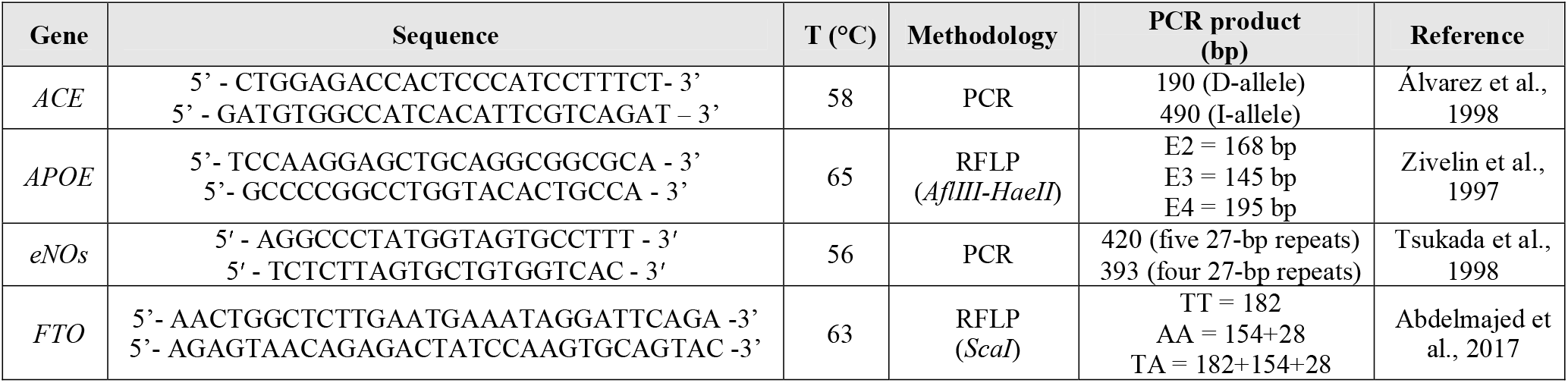
Primers and annealing temperatures for gene polymorphisms analysed in the present study

To verify the genotyping results, 10% of the total sample (n = 110) were also analysed by another investigator. The two analyses showed identical results.

### 2.2 Statistical Analysis

SPSS software statistical package program version 25.0 for Windows (SPSS, Inc., Chicago, USA) was used for all statistical analyses. Pearson’s *χ*^2^ test was conducted for Hardy-Weinberg equilibrium (HWE) and to compare allele or genotype frequencies between different age groups, whereas the ANOVA test was used in order to evaluate possible differences in BMI values between age groups. Finally, multiple regression analysis was also used to evaluate the influence of age on BMI. All P-values were two tailed and the level of statistical significance was set at P<0.05 for all tests.

## 3. RESULTS

In Table 2 the general characteristics of the studied population are reported. A total of 1,100 subjects, 546 males (mean age ± SD: 54.423±21.128, age range: 10-98) and 554 females (mean age ± SD: 57.314±24.038, age range: 10-100) were recruited.

**Table 2.** General characteristics of the studied population.

| Subjects | N (%) | Age, Mean ± S.D.<br>(range) | BMI, Mean ± S.D. |
| --- | --- | --- | --- |
| **Total** | **1100** | **55.879±22.676** | **26.299±4.262** |
| **Sex** |  |  |  |
| Males | 546 (49.64) | 54.423±21.128<br>(10-98) | 26.316±4.114 |
| Females | 554 (50.36) | 57.314±24.038<br>(10-100) | 26.281±4.406 |
| **Age groups** |  |  |  |
| 1-24 | 117 (10.64) | 19.120±4.369 | 24.730±4.307 <sup>a,b</sup> |
| 25-49 | 352 (32.00) | 37.594±6.985 | 26.073±3.797 <sup>c,d</sup> |
| 50-85 | 521 (47.36) | 69.317±9.613 | 27.163±4.359 <sup>e</sup> |
| 86-100 | 110 (10.00) | 89.845±0.279 | 24.598±4.126 |

S.D. = Standard Deviation
<sup>a</sup> $P = 0.015$ with respect to 25-49 age group
<sup>b, c</sup> $P < 0.001$ with respect to 50-85 age group
<sup>d</sup> $P = 0.006$ with respect to 86-100 age group
<sup>e</sup> $P < 0.001$ with respect to 86-100 age group

No significant differences were found between males and females in terms of mean age and BMI values. *Vice versa*, significant differences were found in terms of BMI values between age groups, with exception of 1-24 with respect to 86-100 age groups. This result was confirmed by both the ANOVA analysis (*P* <0.001) and regression analysis (*P* = 0.021) that showed a significant correlation between age and BMI.

In Table 3 the allele and genotype frequencies for all studied genes are reported. All genotype frequencies resulted in accordance with the Hardy-Weinberg law.

**Table 3.** Allele and Genotype Frequencies of four metabolic gene polymorphisms in an Italian sample (n = 1,100)

| Gene polymorphisms | Allele | N | Frequency | Genotype | N | Frequency | HWE<br>*P*-value |
| --- | --- | --- | --- | --- | --- | --- | --- |
| **ACE** | D | 1698 | 0.772 | DD | 643 | 0.585 | 0.110 |
|  | I | 502 | 0.228 | ID | 412 | 0.374 |  |
|  |  |  |  | II | 45 | 0.041 |  |
| **APOE** | E3 | 1855 | 0.843 | E3E3 | 784 | 0.713 | 0.237 |
|  | E4 | 278 | 0.126 | E3E4 | 227 | 0.206 |  |
|  | E2 | 67 | 0.031 | E2E3 | 60 | 0.055 |  |
|  |  |  |  | E4E4 | 22 | 0.020 |  |
|  |  |  |  | E2E4 | 7 | 0.006 |  |
| **FTO** | T | 1417 | 0.644 | TT | 470 | 0.427 | 0.199 |
|  | A | 783 | 0.356 | TA | 477 | 0.434 |  |
|  |  |  |  | AA | 153 | 0.139 |  |
| **eNOs** | 420 | 1842 | 0.837 | 420/420 | 721 | 0.708 | 0.203 |
|  | 393 | 358 | 0.163 | 429/393 | 263 | 0.258 |  |
|  |  |  |  | 393/393 | 35 | 0.034 |  |

HWE = Hardy-Weinberg Equilibrium.
\* The HWE was not calculated because the used genotyping procedure does not consent to discriminate the heterozygotes from homozygotes

Analysing the frequencies of the studied gene polymorphisms (Table 4), we observed a significant decrease of the *APO E4* allele frequency and a significant increase of the *APO E2* allele frequency in the 86-100 age group with respect to 1-24 (*P* = 0.005) and 25-49 (*P* = 0.007) age groups. Similarly, the older age group showed a significant (*P*<0.001) reduction of *eNOs* 393 and *FTO A* alleles with respect to 1-24 and 25-49 age groups. Finally, for ACE, no significant differences in the allele frequencies between age groups were found.

**Table 4.**
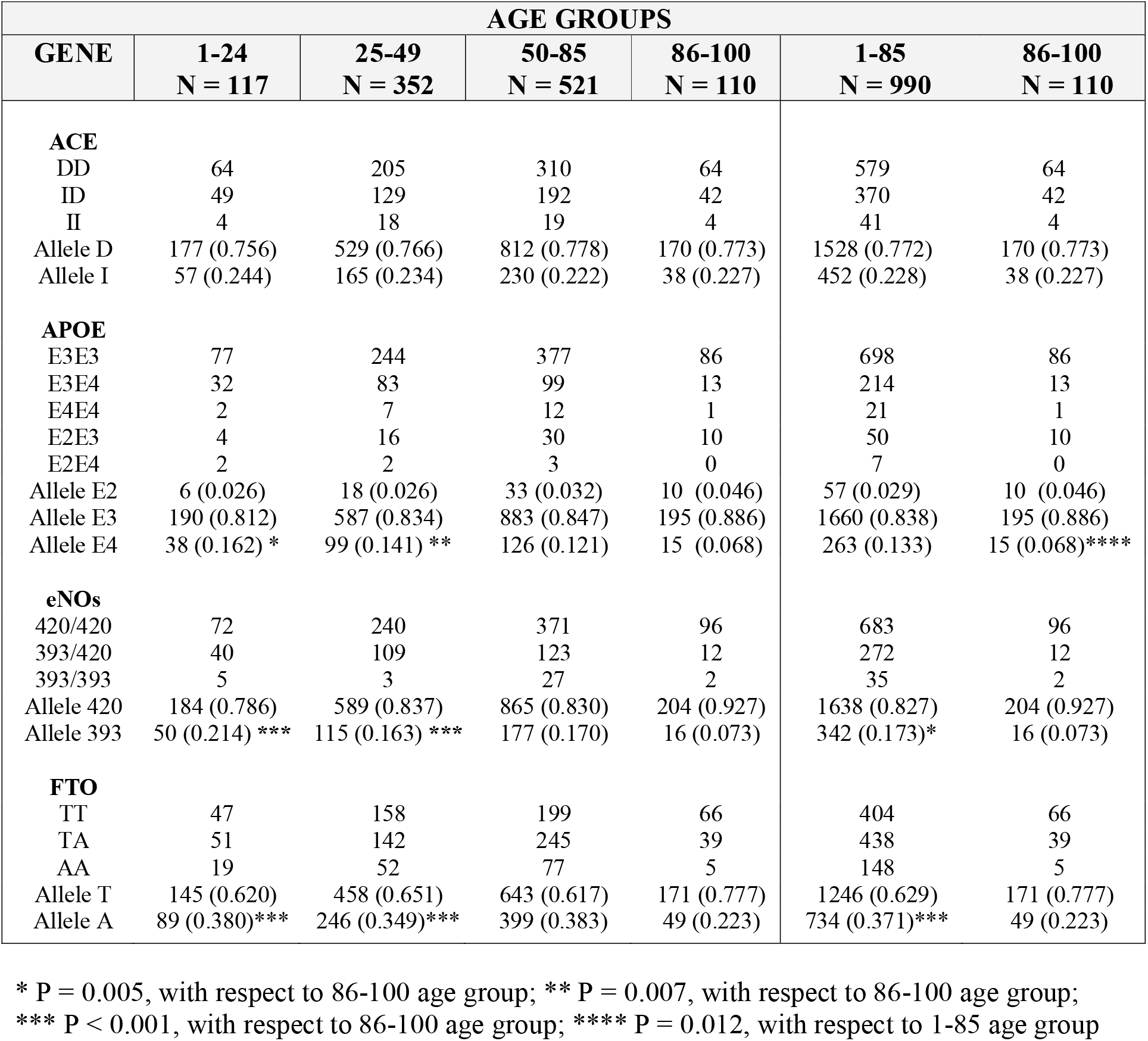
Frequencies of the studied metabolic gene polymorphisms among different age groups

| GENE | AGE GROUPS |  |  |  |  |  |
| --- | --- | --- | --- | --- | --- | --- |
|  | 1-24<br>N = 117 | 25-49<br>N = 352 | 50-85<br>N = 521 | 86-100<br>N = 110 | 1-85<br>N = 990 | 86-100<br>N = 110 |
| **ACE** |  |  |  |  |  |  |
| DD | 64 | 205 | 310 | 64 | 579 | 64 |
| ID | 49 | 129 | 192 | 42 | 370 | 42 |
| II | 4 | 18 | 19 | 4 | 41 | 4 |
| Allele D | 177 (0.756) | 529 (0.766) | 812 (0.778) | 170 (0.773) | 1528 (0.772) | 170 (0.773) |
| Allele I | 57 (0.244) | 165 (0.234) | 230 (0.222) | 38 (0.227) | 452 (0.228) | 38 (0.227) |
| **APOE** |  |  |  |  |  |  |
| E3E3 | 77 | 244 | 377 | 86 | 698 | 86 |
| E3E4 | 32 | 83 | 99 | 13 | 214 | 13 |
| E4E4 | 2 | 7 | 12 | 1 | 21 | 1 |
| E2E3 | 4 | 16 | 30 | 10 | 50 | 10 |
| E2E4 | 2 | 2 | 3 | 0 | 7 | 0 |
| Allele E2 | 6 (0.026) | 18 (0.026) | 33 (0.032) | 10 (0.046) | 57 (0.029) | 10 (0.046) |
| Allele E3 | 190 (0.812) | 587 (0.834) | 883 (0.847) | 195 (0.886) | 1660 (0.838) | 195 (0.886) |
| Allele E4 | 38 (0.162) * | 99 (0.141) ** | 126 (0.121) | 15 (0.068) | 263 (0.133) | 15 (0.068)**** |
| **eNOs** |  |  |  |  |  |  |
| 420/420 | 72 | 240 | 371 | 96 | 683 | 96 |
| 393/420 | 40 | 109 | 123 | 12 | 272 | 12 |
| 393/393 | 5 | 3 | 27 | 2 | 35 | 2 |
| Allele 420 | 184 (0.786) | 589 (0.837) | 865 (0.830) | 204 (0.927) | 1638 (0.827) | 204 (0.927) |
| Allele 393 | 50 (0.214) *** | 115 (0.163) *** | 177 (0.170) | 16 (0.073) | 342 (0.173)* | 16 (0.073) |
| **FTO** |  |  |  |  |  |  |
| TT | 47 | 158 | 199 | 66 | 404 | 66 |
| TA | 51 | 142 | 245 | 39 | 438 | 39 |
| AA | 19 | 52 | 77 | 5 | 148 | 5 |
| Allele T | 145 (0.620) | 458 (0.651) | 643 (0.617) | 171 (0.777) | 1246 (0.629) | 171 (0.777) |
| Allele A | 89 (0.380)*** | 246 (0.349)*** | 399 (0.383) | 49 (0.223) | 734 (0.371)*** | 49 (0.223) |

\* P = 0.005, with respect to 86-100 age group; \*\* P = 0.007, with respect to 86-100 age group;
\*\*\* P < 0.001, with respect to 86-100 age group; \*\*\*\* P = 0.012, with respect to 1-85 age group

A similar trend was also observed subdividing the sample in 1-85 and 86-100 age groups (Table 4). Also in this case, we observed a significant (*P*<0.001) decrease of the *APO E4* and a significant increase of the *APO E2* allele frequencies in the 86-100 age group with respect to 1-85 age group (*P* = 0.012). Moreover, the older age group showed a significant (*P*<0.001) reduction of *eNOS* 393 and *FTO A* alleles with respect to 1-85 age group, whereas for the other three studied gene polymorphisms no significant differences in the allele frequencies were found between age groups.

## 4. DISCUSSION

In humans, exceptional longevity is a complex trait. Studies focused on the determination of genetic features of long-lived individuals, represent one of the main approaches to define the key components of this longevity.

Among the anthropometric parameters associated with longevity, BMI seems to play an important role (Frayling et al., 2007). We reported a significant increase of BMI with age, probably ascribed to a significant increase of this parameter in the 25-49 and 50-84 age groups, as indicated by the non-parametric analysis (Table 2). The lack of significant differences in terms of BMI values between younger and older groups could be explained by the fact that, since in old age there is a change in the balance between fat and muscle mass, BMI becomes less useful in assessing the metabolic health of older adults (Srikanthan and Karlamangla, 2014).

Long-living individuals are characterized by a low incidence of cardiovascular and neurodegenerative diseases (Puca et al., 2012). From genetic point of view, this phenomenon may be due to the presence or absence of specific gene polymorphisms, that confer to the carriers a better ability to counteract cell deterioration during aging (Nijiati et al., 2013). One of the most frequent candidate genes that have been studied for its possible association with longevity were *APOE* gene. In particular, *APOE2* was found to be associated with lower levels of total and LDL cholesterol and with higher levels of HDL cholesterol, while *APOE4* has opposite effects. As consequence, the *APOE4* allele has been linked to atherosclerosis and coronary heart disease and individuals with *E4/E4* genotype are at a higher risk of cardiovascular and Alzheimer’s diseases (Marrzoq et al., 2011). For these reasons, long-lived individuals would be expected to have higher frequencies of E2 allele and lower frequencies of E4 allele with respect to younger age groups. Results of our work seem to confirm these hypotheses, since, in the 86-100 age group, we found a significant decrease of the *APOE4* allele frequency and a significant increase of the *APOE2* allele frequency with respect to younger group.

Differently to *APOE*, data about the association of *ACE I/D* gene polymorphism and longevity are controversy, and results from the published literature provides evidence for a modest positive association between the *ACE* D-allele and *D/D* genotype and exceptional human longevity (for a review see Garatachea et al., 2013). For instance, a published meta-analysis showed that, compared with I/I genotype, the D/D genotype is associated with an increase risk for ischaemic stroke and the D allele with a 14% increased risk of type II diabetes, with respect to the I variant (Zhou et al., 2010). Paradoxically, the same (D) allele which predisposes to CAD has been reported to be more frequent in some cohorts of European centenarians, including from France (Faure-Delanef et al., 1998), United Kingdom (Galinsky et al., 1997), Italy (Seripa et al., 2006), as well as in nonagenarians and centenarians from the Uighur population in China (Rahmutula et al., 2002), suggesting a pleiotropic effect of this gene. However, the association between the D allele and increased longevity has not been replicated in other studies on centenarians from Denmark (Bladbjerg et al., 1999), France (Blanche et al., 2001), Italy (Paolisso et al., 2001), China (Yang et al., 2009) and Spain (Fiuza-Luces, 2011).

Also our data, do not give support for a significant role of the ACE I/D polymorphism on human longevity. These contradictory results could be attributed to different factors, including the ethnic background and the sample size of the cohorts. Indeed, the frequency of the I allele decreases from Northern to Southern Europe, as well as Caucasian and African populations present similar frequencies and differ from those observed in Asian, Amerindian and Polynesian populations (Fiuza-Luces, 2011). Moreover, the sample size limitation is a relative common problem in studies with centenarians owing to the difficulty of gathering cases, being living 100 or more years still a rare phenotype.

*FTO* gene is another longevity-related gene whose common variants were found to be associated to obesity and diabetes (Speakman JR, et al., 2008). In particular, the polymorphism rs9939609, is characterized by the presence of the two common alleles A and T, where the frequency of the minor allele (A) shows a moderate variation in the different European populations, ranging from 0.32 to 0.47 (Osier et al., 2002). In humans, the presence of a copy of the allele A was found to be associated with an average weight increase of 1.2 kg, and the presence of two A alleles with an increase of about 3 kg compared to the TT genotype (Frayling et al., 2007). The increase in adipose tissue in the A allele carriers was found also associated with a atherogenic lipid profile and with higher incidence of myocardial infarction (Doney et al., 2009). Also in this case we can hypothesize a pleiotropic effect of FTO gene that manifests itself with an increased fitness for the allele A carriers in reproductive age and with a lower fitness for the same carriers in old age. Our data confirm this hypothesis, with a frequency of A allele significantly reduced in the older group with respect to the younger.

Finally, we found a significant reduction of the *eNOS 393* allele in the older group with respect to 1-24 and 25-49 age groups. This result seems to confirm the hypotheses of a possible correlation between this gene polymorphism and natural longevity, as postulated by different authors (Nijiati et al., 2013; Puca et al., 2012; Park et al., 2004). It is known that nitric oxide plays a protective role in various important events during atherogenesis, inducing smooth muscle relaxation, inhibition of platelet aggregation, leukocyte adhesion to endothelial cells and preservation of endothelial progenitor cell function (for a review see Puca et al., 2012). However, during aging, there is a progressive misbalance between NO production, which becomes increasingly reduced, and oxidative stress, which increases without a compensatory enhancement of antioxidant defences. As a result, vasodilator function of aged vessels could be compromised, with consequent increase of vascular resistance and impaired perfusion (Forstermann, 2010). In this scenario, *eNOS* may be critically involved in longevity by increasing the deteriorative effects of aging. In particular, the *eNOS 4a* allele and *eNOS 4a/4a* genotype were found to be independent risk factors of myocardial infarction (Park et al., 2004), and their frequencies are expected to be lower in long-lived subjects, as actually observed in our sample.

## 5. CONCLUSIONS

Our results showed an age-related relationship of *APOE, FTO* and *eNOS* gene polymorphisms in a cohort of healthy Italian subjects. Despite the limitation of studies like this, due to the difficulty of finding a substantial number of centenarians and, in general, of long-lived people, it is our opinion that the present work could be relevant for further studies about longevity among Italian and European populations. Finally, it has been estimated that the number of centenarians worldwide will double each decade until 2100 (Nijiati et al., 2013), making these studies increasingly important in the future.

## CONFLICT OF INTEREST

None declared.

